# THE IMPACT OF ARTIFICIAL INTELLIGENCE ON UNIVERSITY STUDENTS’ PERCEPTIONS OF ENVIRONMENTAL CHALLENGES AND SUSTAINABLE SOLUTIONS

**DOI:** 10.1101/2025.07.20.665788

**Authors:** Tien Nhat Huynh, Khuong An Nguyen, Luan Nguyen Trong

**Author notes:** Corresponding Author. Entrepreneurship Department, FPT University, Can Tho Campus, Vietnam. ORCID ID: 0000-0002-3489-1628. [Postal Address: 600 Nguyen Van Cu Street, An Binh Ward, Ninh Kieu District, Can Tho City, 94100, Vietnam].

## Abstract

In the context of Vietnam’s higher education strongly transforming towards digitalization and sustainability, this study aims to analyze the impact of artificial intelligence (AI) on students’ awareness and intention to act for the environment. Based on the Extended Planned Behavior (TPB) model, the research model integrates two input factors: environmental awareness (EA) and the level of interaction with AI in environmental learning (AEEL), affecting three mediating factors: attitude (AT), subjective norm (SN) and perceived behavioral control (PBC), thereby predicting students’ green action intention (GAI). Data collected from 521 students at private university in Vietnam were processed using EFA, CFA and SEM. The results showed that both EA and AEEL had positive effects on AT, SN and PBC (p < 0.001). Among them, AEEL had the strongest influence on SN (β = 0.281) and PBC (β = 0.252). PBC had the strongest influence on GAI (β = 0.312), followed by AT (β = 0.274) and SN (β = 0.267). Bootstrapping mediation analysis confirmed that the mediating variables (AT, SN, PBC) played an important role in linking EA and AEEL with GAI. In addition, cluster analysis showed that female students and senior students had significantly higher levels of AI engagement and action intention. These results suggest that AI not only serves as a learning support tool but also as a catalyst for sustainable behavior in higher education.

## I. Introduction

Under the conditions of increasing and ever more radical climate change, biodiversity loss and ecological pollution actively threatening global quality of life, the challenge of higher education in fostering sustainable consciousness and conduct is more pressing than ever (Chen et al., 2023). In particular, the extraordinary development of artificial intelligence (AI) has been working deep changes not only in educational technology but also in the way in which students cope with, understand, and respond to environmental problems (Vinuesa et al., 2020). AI is increasingly applied to support decision-making, data handling, ecosystem simulation, and environmental risk estimation (Chen et al., 2023). However, there is still limited research on the impact of AI on sustainable action and awareness amongst students – key social agents toward a green future (Zawacki-Richter et al., 2019).

In developing countries like Vietnam, where climate change has very high effects, especially in the Mekong Delta, environmental awareness and green behavior among young people have to be promoted (Do et al., 2025). However, the integration of emerging technologies such as AI into environmental education is yet to be theoretically and empirically investigated (Kabudi, 2022). The majority of the previous literature has emphasized digital literacy or attitude towards technology in education (Zawacki-Richter et al., 2019), but very little literature has described how AI affects students’ attitude and willingness to act in sustainable development (Vinuesa et al., 2020).

Based on this gap, this study builds and tests an integrated theoretical model based on Ajzen’s (1991) Theory of Planned Behavior (TPB), combined with specific factors of AI technology and environmental education (Ajzen, 1991; Kabudi, 2022). Specifically, the proposed theoretical model includes two main input factors, Environmental Awareness and AI Engagement for Environmental Learning, which affect three intermediate components, Attitude, Subjective Norms and Perceived Behavioral Control, which in turn affect students’ Green Action Intention (Ajzen, 1991; Gómez-Ramirez, 2019). This approach not only allows for the analysis of the role of individual cognition but also clarifies the influence of social factors and perceived behavioral control in the context of technology-integrated learning (Ajzen, 1991).

The inclusion of the factor "AI Engagement for Environmental Learning" is a prominent new feature of the model, reflecting the topicality and convergence between technology and environmental education (Ncube & Ngulube, 2024). This factor is understood as the extent to which students use, interact with, and learn through AI tools to better understand environmental issues, thereby forming a cognitive foundation and the ability to respond positively (Do et al., 2025). Tools such as learning support chatbots, environmental data analysis platforms, or pollution forecasting applications using AI are increasingly popular in universities in Vietnam (Do et al., 2025), opening up new learning spaces for the student generation (Zawacki-Richter et al., 2019).

At the same time, the environmental awareness factor plays a fundamental role in the model. It has been established through several studies that increased attention towards issues such as climate change, resource shortage, or plastic pollution is positively correlated with green action-supporting attitudes and green action intentions to participate in environmental protection activities (Kollmuss & Agyeman, 2002). However, the extent to which this perception is influenced depends on the mediating factors of social norms and perceived ability to perform the behavior, more so in the academic setting where students are motivated by teachers, peers, and the education system (Lee, 2009).

Environmental action attitudes are the first important factor to consider. This is what captures whether or not students have positive or negative attitudes towards performing sustainable actions, such as saving energy, reducing waste, or participating in neighborhood green projects (Nguyen Van & Le Hoang, 2024). With high environmental awareness among students and being prompted to use AI for exploring sustainable alternatives, positive attitudes can be fostered (Do et al., 2025). Subjective norms are the second component, reflecting perceptions of social expectations – such as peers, instructors, or the academic community – for green behavior (Lee, 2009). The use of AI may increase perceptions of consensus or positive pressure from the surrounding environment, thereby strengthening social norms toward sustainable behavior (Thanh & Toan, 2023). Finally, perceived behavioral control refers to students’ feelings about their ability to perform green behavior based on resources and support they have, including technological capabilities and institutional support (Ajzen, 1991).

From the three mediating components above, the model hypothesizes that these factors will directly impact green action intentions – an important indicator predicting the likelihood that students will engage in sustainable behaviors in the present and future (Ajzen, 1991). Based on the specific hypotheses (H1a–H5), the study tests the relationships between the factors using quantitative methods and SEM structural analysis (Nguyen Van & Le Hoang, 2024).

The case study is Can Tho Private University – a pioneer in applying AI in teaching and research. The survey here helps ensure practicality and representation of the technology education context in the Mekong Delta region, which is facing serious environmental challenges such as salinity intrusion, land subsidence, and groundwater pollution (Trong Nguyen & Tran, 2024). How students here perceive and are willing to act for the environment with the support of AI is a question of profound scientific and practical value (Do et al., 2025).

This study contributes in three main aspects. First, in terms of theory, the study extends the Theory of Planned Behavior by integrating modern technological elements such as AI, creating a multidimensional analytical framework between technology, education and environmental behavior (Ajzen, 1991). Second, in terms of experimentation, the study provides real data from students in the context of digital transformation of education in Vietnam, where both AI and environmental education are in the process of strong development but have not been closely integrated (Ngo et al., 2020). Third, in terms of policy, the research results can help education managers design more effective AI training programs and policies, aiming to form ecologically responsible digital citizens - an urgent requirement in the current era of digital transformation and climate crisis (Chen et al., 2023).

In summary, this study not only expands the understanding of the impact of new technologies on environmental awareness and behavior among young people, but also provides a scientific basis for integrating artificial intelligence into sustainable education strategies (Vinuesa et al., 2020).

The results are expected to contribute to the interdisciplinary theoretical framework between technology – education – sustainable development, and help universities in Vietnam shape green digital transformation strategies that are suitable for the current generation of students (Do et al., 2025). Based on the theoretical and practical foundations mentioned above, the next section will present the theoretical foundation system as the basis for building a research model, focusing on key concepts such as environmental behavior, technological awareness and the participation of artificial intelligence in sustainable education.

## II. Literature Review

### 1. Theoretical background

#### Feasibility of the TPB model in environmental behavior research

The Theory of Planned Behavior (TPB) developed by (Ajzen, 1991) has been widely recognized as an effective analytical framework for individual behaviors in many fields, including environmentally friendly behavior (Ajzen, 1991; Tsai & Tan, 2022). According to TPB, attitudes – that is, individuals’ positive or negative views about a particular behavior – play a central role in shaping behavioral intentions, thereby accurately predicting green behavioral patterns such as recycling, energy saving, and sustainable consumption (J.-W. Wang et al., 2021). In addition, subjective norms reflect perceived social pressure from relatives, friends, and the academic community; Several meta-analyses have confirmed that this factor contributes significantly to the intention to adopt green behavior, although the extent of its influence may vary according to cultural context (Bamberg & Möser, 2007). Finally, perceived behavioral control captures an individual’s perception of the ability and resources to perform the desired behavior; this factor not only influences intention but also directly influences actual behavior when sufficient support conditions are present (Li et al., 2023).

The practical value of the TPB is reinforced by several meta-analyses showing that this model can explain an average of 39% of the variance in behavioral intention and about 27% in actual environmental behavior (Armitage & Conner, 2001). Furthermore, studies using structural analysis methods such as PLS-SEM in university contexts have shown that attitude is the strongest predictor, followed by social norms and behavioral control (Phang & Ilham, 2023). A recent study at the University of Malaya confirmed that all three factors in the TPB significantly influence students’ green behavior, especially in technology-supported environmental education programs (Phang & Ilham, 2023). From an applied perspective, the TPB provides a clear theoretical framework to identify and classify psychological variables (attitude, norms, and control) that influence green behavior, while also providing a suitable foundation for integrating more modern factors such as AI engagement in the context of digital education in developing countries such as Vietnam (NGUYEN et al., 2024). From the foundation of the TPB model, the next section will clarify the role of AI Engagement in environmental education, especially in the context of higher education in developing countries like Vietnam.

#### AI Engagement and Environmental Learning Dynamics

In the context of modern education, AI engagement refers to the extent to which students use, interact with, and experience AI tools during their learning process (Sajja et al., 2025). A recent study by OceanChat found that interactive chatbots in the form of marine creatures (such as whales) have the potential to enhance students’ awareness and sustainable behavioral intentions through personalized dialogue (Pataranutaporn et al., 2025). The results showed that the real-time dialogue model with the character was more inspiring than traditional learning (Pataranutaporn et al., 2025). This demonstrates that AI Engagement is not just about using but also deeply interacting with environmental content (Phang & Ilham, 2023).

AI tools such as AI-Enabled Intelligent Assistant (AIIA) have been developed to support personalized education by providing feedback, querying knowledge, and adjusting learning paths (Sajja et al., 2025). Studies in the field of environmental engineering have shown that students find it easier and more convenient to use AI than learning through a traditional teacher or tutor (Sajja et al., 2025). However, the authors also note that clear policy and ethical support are needed to ensure responsible learning (Sajja et al., 2025).

In environmental education, AI-enabled teaching systems have been shown to improve knowledge, attitudes, and green action intentions, which are important (Huang, 2018). Research from an AI-VR system in the Philippines shows that dynamic simulations (such as sea level rise, deforestation) help students gain a deeper understanding of ecosystems and motivate them to engage in green actions (Huang, 2018). In addition, AI promotes critical thinking and sustainable values when students are deeply engaged with real-world situations through technology (Hajj-Hassan et al., 2024).

In addition, research conducted by indicates that AI technologies such as ChatGPT and Generative AI allow for the optimization of learning resources, increase accessibility, and promote environmental sustainability in the university environment (Nikolopoulou, 2025). They suggest that AI policies in the university environment should be developed to promote responsible active learning experiences (Okulich-Kazarin et al., 2023).

In summary, AI Engagement plays a central role in developing environmental learning motivation by providing interactive, personalized experiences and real-time feedback. When properly integrated in higher education environments, AI not only supports knowledge but also creates a foundation for positive attitudes, perceived control and subjective norms towards green behavior (Nikolopoulou, 2025). This explains why the extended TPB model with AI Engagement variable is appropriate and necessary in research in Vietnam (Thanh Binh et al., 2025). Building on this foundation, the next section discusses how emerging technologies like AI can be integrated into TPB to enhance its relevance.

#### Extending TPB with AI-related factors

In the context of studying green behavior supported by AI, many scholars have proposed to extend the traditional TPB model by adding digital technology factors such as “perceived usefulness” and “perceived ease of use” from TAM, as well as “trust in AI” and “AI attributes” to enhance the explanatory power of behavior (H.-H. Yang & Su, 2017). For example, research combining TPB and TAM shows that perceived usefulness of AI has a direct impact on attitudes and intentions to use AI in an environmental context, thereby indirectly promoting environmentally friendly behavior (Y. Yang & Kim, 2024).

Similarly, perceived ease of use has been shown to positively influence behavioral control and attitudes through PLS-SEM systems, indicating that when students perceive AI as accessible, they feel more confident in applying it to the learning environment (Liao, 2025).

Furthermore, trust in AI is increasingly viewed as a moderator variable that influences attitudes and subjective norms in the extended TPB model. Recent research from IEEE indicates that AI attributes such as “perceived intelligence,” “perceived anthropomorphism,” and “perceived animacy” directly influence users’ beliefs and attitudes, thereby enhancing their intention to engage in socially sustainable activities (Choung et al., 2023). This is consistent with the results from AI-enabling energy-saving support systems, where perceived behavioral control and perceived usefulness were found to be the most important factors driving intention to use (Y. Yang & Kim, 2024).

It is important to note that many models integrating AI attributes in the TPB also show that digital communication and environmental information provided by AI can reshape social norms through online learning networks (Popkova et al., 2022). Accordingly, when AI provides data on energy efficiency, carbon sequestration, or community contact in real time, individuals who access this information undergo a process of digital socialization, which can change subjective norms and promote green behavior (Popkova et al., 2022; Sarmento & Loureiro, 2021).

Finally, in-depth studies using SEM-ANN techniques to analyze the extended TPB model show that intention to use AI as a mediator plays an essential role in creating green behavior (Soomro et al., 2025). The model indicated that intention to use AI is an important mediator between variables such as perceived usefulness, ease of use, and AI trust with green behavior (Soomro et al., 2025). This is strong evidence that extending the TPB with AI-related factors is not only feasible but also necessary to model green behavior in the current digital context (Armitage & Conner, 2001).

In summary, the integration of AI technology variables (usefulness, ease of use, system trust, and AI attributes into the TPB model has been verified by the latest quantitative analysis and shows a stronger explanatory power for green behavior among students and digital homemakers (Phang & Ilham, 2023). This is a suitable approach to apply research in Vietnam, where the extended TPB model with AI can explain the mechanisms affecting students’ attitudes, norms, behavioral control, and ultimately sustainable behavior (NGUYEN et al., 2024). Given this integration of TPB and AI-related factors, it is essential to clearly define the research variables involved.

### 2. Review of the literature

#### Environmental Awareness

Environmental awareness is defined as the level of understanding, concern, and awareness of individuals about environmental issues such as climate change, pollution, ecosystem degradation, and personal responsibility for action (Kollmuss & Agyeman, 2002). It is a fundamental cognitive factor that facilitates the formation of positive attitudes toward green behavior (Li et al., 2023). Empirical studies show that people with high environmental awareness tend to make environmentally friendly decisions, from product selection to participation in community activities (Lee, 2009). In higher education, increasing environmental awareness is a core goal of the sustainable development program (Xiao et al., 2024).

#### AI Engagement for Environmental Learning

This variable measures the extent to which students use, interact with, and learn through AI tools such as chatbots, learning support systems, digital environmental simulations, or climate data analysis applications (Sajja et al., 2025). AI Engagement not only reflects digital behavior, but is also an indicator of the level of active engagement in innovative learning environments (Sajja et al., 2025). Research shows that high levels of engagement with AI help students develop critical thinking skills and a deeper understanding of sustainable solutions (Pataranutaporn et al., 2025). This factor is considered a modern variable that needs to be integrated into the TPB model to reflect the current digital education context (Ajzen, 1991; Nopas, 2025).

#### Attitude

Attitude refers to how far students evaluate environmental behaviors such as conserving water, recycling, planting trees, or reducing emissions as positive or negative (Ajzen, 1991). Positive attitude is established when the students find green behavior useful, beneficial, and complementary to their personal values. Attitude has also been identified to be the strongest predictor of behavioral intention in the majority of TPB models (Armitage & Conner, 2001). Green dispositions in this research could be influenced by environmental knowledge and exposure to learning using AI technology (Lee, 2009).

#### Subjective Norms

Subjective norms are students’ perceptions regarding what other people, including friends, teachers, family, or the academic community, expect of them regarding green behavior (Bamberg & Möser, 2007). The more students perceive that people close to them support environmentally friendly behavior, the more they are motivated to do so (Kollmuss & Agyeman, 2002). Social norms play a deep significance in the education settings, where group learning and social discussion become the focal points (Li et al., 2023). Additionally, AI technology indirectly affects social norms through content recommendation and internet group interactions (Kabudi, 2022).

#### Perceived Behavioral Control

Perceived behavioral control measures the extent to which students feel they have the ability, knowledge, tools, and support to engage in green behavior (Ajzen, 1991). When students believe that they have easy access to resources such as technology, curriculum, or time, behavioral intentions increase significantly (Xiao et al., 2024). In the context of digital transformation, the perception of the ability to control AI tools and digital learning environments is also an inseparable part of perceived control (Nopas, 2025). Therefore, this factor is a bridge between personal capacity and external support conditions (Li et al., 2023).

#### Green Action Intention

Green action intention reflects the level of commitment and willingness of students to perform environmentally beneficial behaviors in the near future, such as changing consumption habits, participating in volunteering, or applying sustainable solutions in practice (S. Wang et al., 2025). This is the central dependent variable in TPB, formed by three mediating factors: attitude, subjective norms, and perceived control (Armitage & Conner, 2001). Many studies confirm that intention is a necessary premise to lead to actual behavior, and is a reliable indicator in measuring the effectiveness of environmental education interventions (Ajzen, 1991;

Patel et al., 2018). In this study, green action intention also plays a role in reflecting the level of transformation of awareness and technology into action. Although the variables have been conceptually defined, a critical examination of the research gap is necessary to justify the model’s contribution.

#### H1a

There is a positive relationship between environmental awareness and students’ attitude toward green action.

#### H1b

There is a positive relationship between environmental awareness and students’ subjective norms.

#### H1c

There is a positive relationship between environmental awareness and students’ perceived behavioral control.

#### H2a

There is a positive relationship between AI engagement for environmental learning and students’ attitude toward green action.

#### H2b

There is a positive relationship between AI engagement for environmental learning and students’ subjective norms.

#### H2c

There is a positive relationship between AI engagement for environmental learning and students’ perceived behavioral control.

#### H3

There is a positive relationship between students’ attitude toward green action and their green action intention.

#### H4

There is a positive relationship between students’ subjective norms and their green action intention.

#### H5

There is a positive relationship between students’ perceived behavioral control and their green action intention.

**Fig. 1.**
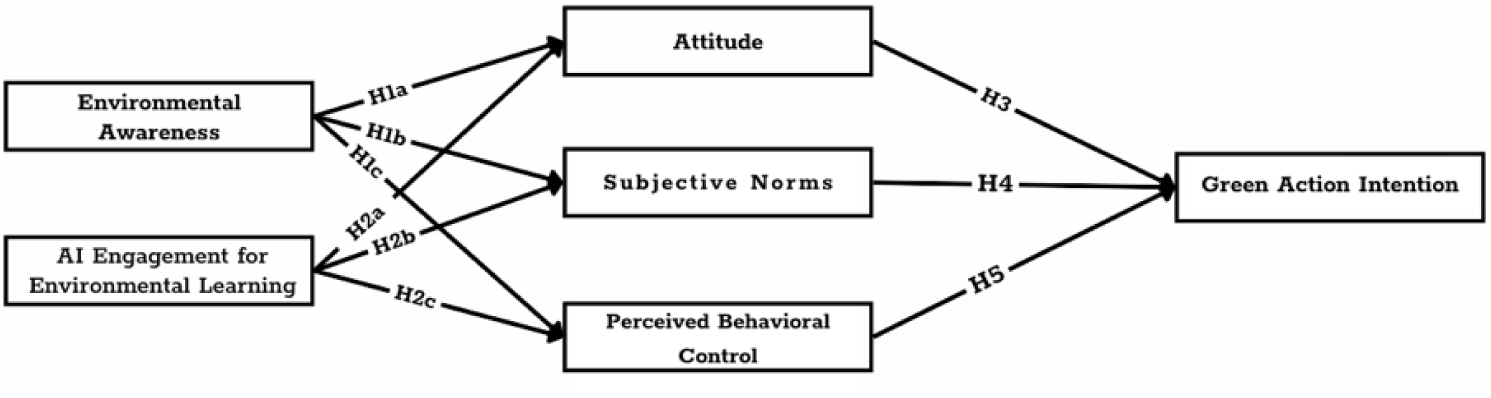
Conceptual framework

#### Research Gap

Although the TPB model has been extensively used to predict environmental behavior in a wide range of contexts, most recent studies still focus on the traditional psychological determinants of attitudes, subjective norms, and perceived behavioral control (Armitage & Conner, 2001; Bamberg & Möser, 2007). Meanwhile, the incorporation of artificial intelligence (AI) into education has not been wholly accomplished in behavioral models such as the TPB, especially in developing countries’ environmental education (Hajj-Hassan et al., 2024; Pataranutaporn et al., 2025).

In addition, very few studies have considered the role of AI Engagement in measuring the extent to which students interact with AI tools in environmental learning – as a fundamental factor influencing the psychological components in the TPB (Hajj-Hassan et al., 2024; Huang, 2018). In Vietnam, there has been almost no quantitative research combining the extended TPB with AI technology factors to predict students’ green action intentions in the context of digital transformation (Thanh Binh et al., 2025). This gap poses an urgent need for integrated theoretical models that both inherit the traditional TPB structure and reflect the modern practice of environmental education in the AI era. Based on the identified gaps, the following section outlines the proposed research model and hypotheses.

## III. Methodology of Research

### 1. Sample size

The present study was performed on university students, who were studying in one of the private universities at Can Tho city, Vietnam. The study on persons’ attitudes towards environmental problems and willingness to act green in using AI for learning. The general size of the University student population may be significant but only students from various faculties and academic years were focused on to guarantee sufficient representation and research stability.

According to Hair et al. (2013), the ideal sample size for Applying EFA is supposed to be a 5:1 ratio of Observations to Variables or 10:1 Observations to Variables. Because the questionnaire consisted of 30 items belonging to six factors, the minimum number of respondents AVE must be 30*5 = 150 cases. To enhance statistical power, the study aimed to exceed this threshold.

In all, 521 valid responses were received. Following thorough screening processes and removal of incompletely fulfiled or invalid responses, 521 responses were used for analysis. This sample size is sufficient for SEM analysis, and it provides robust parameter estimation.

### 2. Sampling Methods

Convenience sampling was used in the study to collect students who were readily available and self-selected. The survey was distributed to those with an online platform, student connection, or classroom visit. This method was selected because it allowed us to collect a substantial and diverse number of responses in a short amount of time, and at the same time encompass a broad range of academic majors and year in school.

Participation was entirely voluntary and all respondents were told about the confidential nature of the research and that it was being conducted in an academic context. We applied a strict filtering to the raw dataset retrieved. The resulting final sample of 521 valid responses accounted for 89% usable data and met the requirements for both exploratory and confirmatory analysis.

### 3. Data Analysis Method

The data were processed by SPSS 26 and AMOS 24 with the method of multistep analysis. Descriptive statistics were first used to profile demographic traits and students’ exposure to AI and ecological consciousness. Second, reliability was examined by using Cronbach’s Alpha, which showed that all constructs were higher than 0.84, indicating that the internal consistency was high. Third, EFA based on principal axis factoring and Promax rotation verified the underlying factor structure with a KMO of 0.939 and six extracted factors. Fourth, the measurement model was validated by Confirmatory Factor Analysis (CFA), which provided good fit indices (χ²/df = 1.179, RMSEA = 0.019, CFI = 0.990, GFI = 0.946) and confirmed convergent and discriminant validity. Fifth, hypothesis tests were conducted via Structural Equation Modeling (SEM) indicating significant direct and indirect effects according to the extended TPB model. Group differences based on the demographic variables were also tested with Independent Sample T-tests and One-way ANOVA.

### 4. The Need for Research

The private University student body should see this research immediately because of the university commitment to practice-oriented training. For this orientation, the OJT training paradigm is appropriate as it enables students to put their knowledge and learning skills into practice. To increase their competitiveness in the job market and gain more real-world experience, a large number of students at private University wish to participate in OJT. Companies need to fill positions with experienced private University students, and OJT helps students gain real-world experience, which satisfies the needs of hiring managers.

The importance of this research is demonstrated by its evaluation of the impact of on-the-job training (OJT) on students at private University’s work environment, engagement, learning motivation, career growth, and job satisfaction. Determine what aspects of the OJT program, the corporate environment, and the personal traits of the students all have an impact on how effective the program is. Provide private University students with efficient results from the OJT model outcomes that meet their demands as well as the needs of businesses and the university.

## IV. Results

### 1. Descriptive Statistics

In order to ensure the reliability and generalizability of the sample data for SEM analysis, descriptive statistics were carried out to examine the demographic profiles of the participants. There were 521 valid responses collected from private university in Vietnam after survey. Malhotra (2020) indicates a sample size larger than 200 is sufficient in SEM studies, especially when the model contains multiple latent constructs and observed variables. Thus the yielded sample fulfilled necessary statistical assumption for further analysis.

Regarding gender, a representative of the sample were represented by 58.3% male and 41.7% female participants. Based on academic level, 22.3% of respondents were first-year students, 23.2% were in the second year of study, 30.1% in the third year, and 24.4% in the fourth year. This coverage indicates that the student participants represent every level of the academic hierarchy. By field of study 35.7% of respondents were pursuing business degrees, 26.3% degrees in the social sciences, 20.1% degrees in technology or engineering, and 17.9% other disciplines, underscoring the interdisciplinary nature of the research topic.

These statistics demonstrate that the sample is diverse and thus generalizable. Also, the variation in academic year and major type will support meaningful subgroup analyses (T-test and ANOVA) of differences in AI engagement patterns and environmental perception based on demographic characteristics.

**Table 1.** Demographic Profile of Respondents.

| Variable | Category | Frequency (n) | Percentage (%) |
| --- | --- | --- | --- |
| Gender | Male | 304 | 58.3 |
|  | Female | 217 | 41.7 |
| Academic Year | First Year | 116 | 22.3 |
|  | Second Year | 121 | 23.2 |
|  | Third Year | 157 | 30.1 |
|  | Fourth Year | 127 | 24.4 |
| Academic Major | Business | 186 | 35.7 |
|  | Social Sciences | 137 | 26.3 |
|  | Technology & Engineering | 105 | 20.1 |
|  | Others | 93 | 17.9 |

### 2. Reliability Analysis – Cronbach’s Alpha

The validity of the measurement scales was tested using Cronbach’s Alpha coefficients to measure the internal consistency of the scales. According to Hair et al. (2013) and Nunnally & Bernstein (1994) a Cronbach’s Alpha (α) above 0.70 is indicative of acceptable reliability, and above 0.80 as high reliability. Additionally, in order to confirm the contribution of each of the items to the overall scale reliability, its Corrected Item–Total Correlation (CITC) has to be higher than 0.30 (Churchill, 1979).

The results showed that all of the six constructs fulfilled the reliability requirements. In particular, EA (α =.861) and all CITC were greater than 0.45, indicating a scale with internal consistency. AI Engagement for Environmental Learning (AEEL) also demonstrated good internal consistency (a = 0.870). Values of alpha from 0.847 to 0.865 were obtained for the AT (Attitude Toward Green Action), SN (Subjective Norms), and PBC (Perceived Behavioral Control) scales. Lastly, GAI has the highest internal reliability of all constructs, α = 0.880.

**Table 2.** Reliability Analysis of Constructs (Cronbach’s Alpha)

| Construct | Number of Items | Cronbach’s Alpha | Reliability Level |
| --- | --- | --- | --- |
| Environmental Awareness (EA) | 5 | 0.861 | High |
| AI Engagement for Env. Learning | 5 | 0.870 | High |
| Attitude Toward Green Action (AT) | 5 | 0.847 | High |
| Subjective Norms (SN) | 5 | 0.857 | High |
| Perceived Behavioral Control (PBC) | 5 | 0.865 | High |
| Green Action Intention (GAI) | 5 | 0.880 | Very High |

These results indicate that all constructs are internally consistent and qualify for the second analysis of the factors. No item was excluded at this phase supporting the model’s theoretical underpinning and corroborating our research aim of confirming the potential of AI engagement and environmental attitudes for shaping green behavior intentions.

### 3. Exploratory Factor Analysis (EFA)

We performed an EFA to determine the structural design of the observed variables and for the purpose of testing construct validity before CFA was employed. This method is especially valuable when checking or revising scales in new culture or new organization (Hair et al., 2013). The analysis was conducted using Principal Axis Factoring with Promax rotation in order to allow the constructs to correlate which is consistent with behavioral and attitudinal research.

#### a. Sampling Adequacy and Factorability

To evaluate whether the dataset was suitable for factor analysis, two preliminary tests were conducted. The Kaiser-Meyer-Olkin (KMO) measure of sampling adequacy was 0.939, “marvelous” according to the criterion recommended by Kaiser (1974) and significantly above the accepted value of 0.60 as per Hair et al. (2013). The Bartlett’s Test of Sphericity was found to be extremely significant ( χ² = 7230.225, df = 435, p <. 001), suggesting that the matrix of correlations was different from an identity matrix and that they contained enough correlations between the variables to allow for factor extraction.

**Table 3.** KMO and Bartlett’s Test of Sphericity.

| Indicator | Value |
| --- | --- |
| KMO Measure of Sampling Adequacy | 0.939 |
| Bartlett’s Test – Chi-Square Approx. | 7230.225 |
| Degrees of Freedom (df) | 435 |
| Significance (Sig.) | 0.000 |

#### b. Factor Extraction and Total Variance Explained

The extraction yielded six factors with eigenvalues greater than 1.0, cumulatively explaining 63.1% of the total variance. According to Hair et al. (2013), a cumulative variance above 50% is acceptable for establishing construct validity in social sciences research. Each factor represents one of the six theoretical constructs proposed in the extended Theory of Planned Behavior, affirming the appropriateness of the model in the Vietnamese educational context.

**Table 4.**
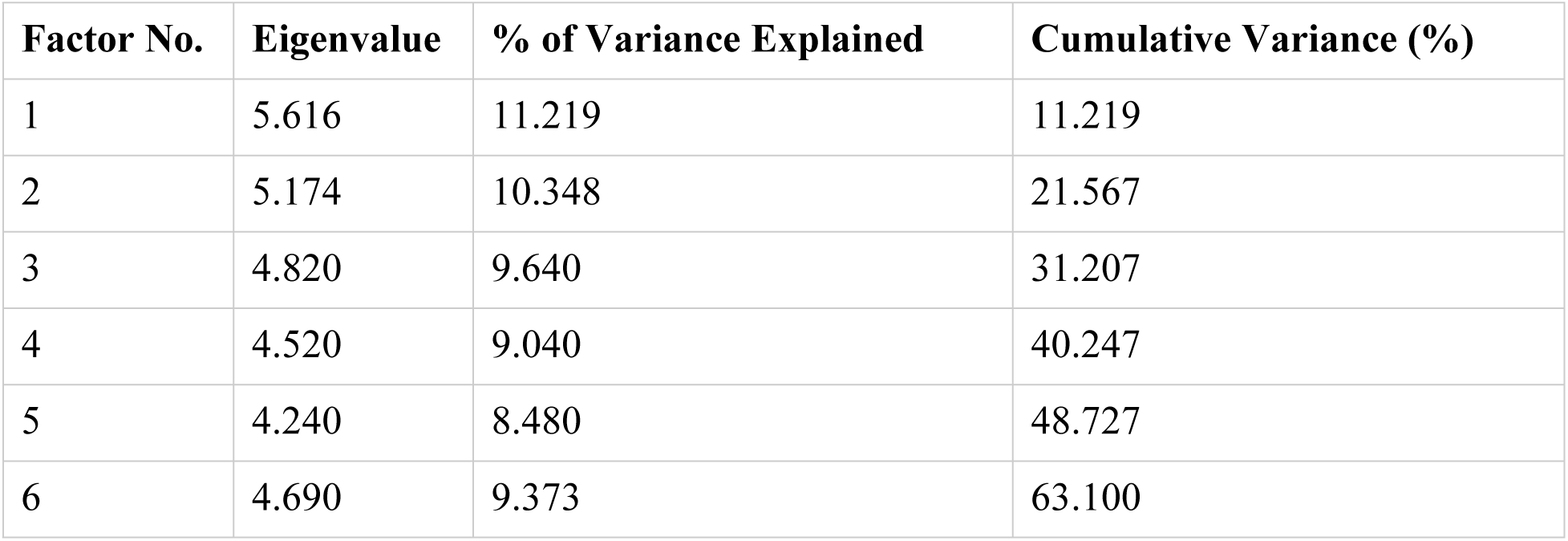
Total Variance Explained by Extracted Factors.

| Factor No. | Eigenvalue | % of Variance Explained | Cumulative Variance (%) |
| --- | --- | --- | --- |
| 1 | 5.616 | 11.219 | 11.219 |
| 2 | 5.174 | 10.348 | 21.567 |
| 3 | 4.820 | 9.640 | 31.207 |
| 4 | 4.520 | 9.040 | 40.247 |
| 5 | 4.240 | 8.480 | 48.727 |
| 6 | 4.690 | 9.373 | 63.100 |

#### c. Rotated Factor Matrix and Factor Loadings

The rotated factor matrix indicated that all items loaded significantly on their intended factors, with factor loadings exceeding 0.50 and cross-loadings remaining below 0.30. These results meet the thresholds recommended by Field (2013) and Costello & Osborne (2005), thus confirming both convergent and discriminant validity of the constructs. The clear and distinct factor structure aligns well with the theoretical framework and supports the scale’s construct integrity for subsequent CFA.

**Table 5.** Factor Loading and Cross-loading Assessment.

|  | Factor |  |  |  |  |  |
| --- | --- | --- | --- | --- | --- | --- |
|  | 1 | 2 | 3 | 4 | 5 | 6 |
| **AT5** | .759 |  |  |  |  |  |
| **AT1** | .749 |  |  |  |  |  |
| **AT4** | .730 |  |  |  |  |  |
| **AT3** | .711 |  |  |  |  |  |
| **AT2** | .681 |  |  |  |  |  |
| **SN5** |  | .782 |  |  |  |  |
| **SN2** |  | .748 |  |  |  |  |
| **SN1** |  | .727 |  |  |  |  |
| **SN4** |  | .703 |  |  |  |  |
| **SN3** |  | .635 |  |  |  |  |
| **EA2** |  |  | .752 |  |  |  |
| **EA3** |  |  | .749 |  |  |  |
| **EA5** |  |  | .712 |  |  |  |
| **EA4** |  |  | .710 |  |  |  |
| **EA1** |  |  | .659 |  |  |  |
| **PBC2** |  |  |  | .782 |  |  |
| **PBC5** |  |  |  | .758 |  |  |
| **PBC1** |  |  |  | .751 |  |  |
| **PBC4** |  |  |  | .699 |  |  |
| **PBC3** |  |  |  | .656 |  |  |
| **GAI4** |  |  |  |  | .767 |  |
| **GAI3** |  |  |  |  | .725 |  |
| **GAI5** |  |  |  |  | .719 |  |
| **GAI2** |  |  |  |  | .719 |  |
| **GAI1** |  |  |  |  | .650 |  |
| **AEEL1** |  |  |  |  |  | .755 |
| **AEEL5** |  |  |  |  |  | .754 |
| **AEEL2** |  |  |  |  |  | .704 |
| **AEEL3** |  |  |  |  |  | .696 |
| **AEEL4** |  |  |  |  |  | .652 |
| Extraction Method: Principal Axis Factoring.<br>Rotation Method: Promax with Kaiser Normalization. |  |  |  |  |  |  |

### 4. Confirmatory Factor Analysis (CFA)

Confirmatory Factor Analysis (CFA) was conducted to validate the measurement model and to confirm the relationships between observed indicators and their corresponding latent constructs. This procedure ensures the unidimensionality, reliability, and validity of the constructs before proceeding to structural model testing (Byrne, 2016; Hair et al., 2013).

#### a. Model Fit Evaluation

The model fit assessment was evaluated by various indices. The value for the chi-square by degrees of freedom ratio (CMIN/df) was 1.179, lower than the criterion of 3.0 recommended, which indicates a parsimonious model (Kline, 2016). The Root Mean Square Error of Approximation (RMSEA) was 0.019, below the 0.08 criterion, indicating a good model fit (Hair et al., 2013). Furthermore, the Comparative Fit Index (CFI = 0.990), Tucker–Lewis Index (TLI = 0.989) and GFI = 0.946 surpassed the recommended critical value of 0.90, suggesting that the model has considerable empirical validity.

**Table 6.** Model Fit Indices.

| **Fit Index** | **Value** |
| --- | --- |
| CMIN/df | 1.179 |
| RMSEA | 0.019 |
| CFI | 0.990 |
| GFI | 0.946 |
| TLI | 0.989 |

Taken together, these findings provide empirical support for the adequacy of the measurement model, implying no significant difference between the proposed model and the empirical data.

#### b. Convergent Validity and Reliability

Standardized factor loadings, Average Variance Extracted (AVE), and Composite Reliability (CR) were examined to assess convergent validity. All loadings were greater than 0.65 and were all significant at 0.001, providing evidence that the factor loads the items highly (Hair et al., 2013).

The AVE values for all six constructs varied from 0.514 to 0.555, whereby all sizes are over 0.50 that demonstrates more than 50% of the variance in the indicators is from the latent construct. The Composite Reliability ranged between 0.841 and 0.862, which exceeded the cutoff value of 0.70 indicating the internal consistency of the measurement model.

**Table 7.** Convergent Validity and Reliability.

| Construct | Standardized Loadings | AVE | CR |
| --- | --- | --- | --- |
| Environmental Awareness (EA) | 0.728 – 0.810 | 0.525 | 0.847 |
| AI Engagement (AEEL) | 0.700 – 0.787 | 0.514 | 0.841 |
| Attitude Toward Green Action (AT) | 0.716 – 0.820 | 0.545 | 0.857 |
| Subjective Norms (SN) | 0.729 – 0.792 | 0.538 | 0.853 |
| Perceived Behavioral Control (PBC) | 0.731 – 0.798 | 0.555 | 0.862 |
| Green Action Intention (GAI) | 0.752 – 0.844 | 0.522 | 0.845 |

These findings support that each construct in the model is both reliable and exhibits satisfactory convergent validity.

#### c. Discriminant Validity

Discriminant validity was tested with the Fornell–Larcker criterion. This rule implies that the square root of the AVE for each construct must be higher than its correlation with any other constructs. Results All constructs satisfy this condition.

Moreover, the MSV of all constructs is less than AVE value for each construct which further approves the discriminant validity.

**Table 8.** Discriminant Validity – Fornell–Larcker Criterion.

| Construct | $\sqrt{\text{AVE}}$ | AT | GAI | EA | AEEL | SN | PBC |
| --- | --- | --- | --- | --- | --- | --- | --- |
| AT | 0.738 | 0.738 |  |  |  |  |  |
| **GAI** | 0.723 | 0.532 | 0.723 |  |  |  |  |
| **EA** | 0.724 | 0.506 | 0.534 | 0.724 |  |  |  |
| **AEEL** | 0.717 | 0.519 | 0.489 | 0.499 | 0.717 |  |  |
| **SN** | 0.733 | 0.549 | 0.532 | 0.513 | 0.521 | 0.733 |  |
| **PBC** | 0.745 | 0.564 | 0.572 | 0.560 | 0.530 | 0.559 | 0.745 |

### 5. Structural Equation Modeling (SEM)

The purpose of this section is to test the structural relationships among latent variables as hypothesized in the extended Theory of Planned Behavior (TPB). Structural Equation Modeling (SEM) was applied using the Maximum Likelihood Estimation method in AMOS 24.

**Fig. 2.**
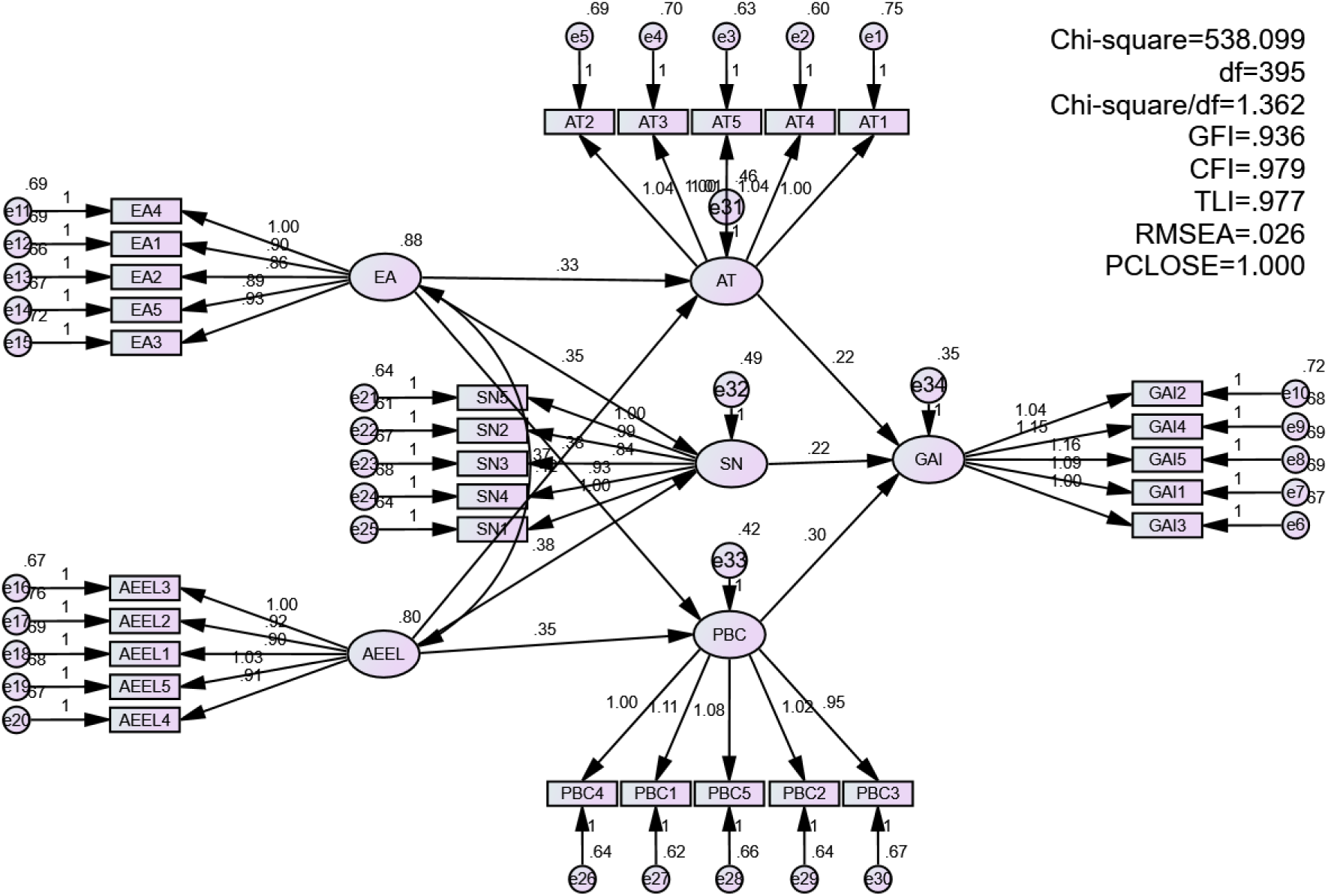
SEM model

#### a. Model Fit Indices

The fit of the structural model was good. The CMIN/df value was 1.235, at the recommended value lower than 3.0 by (Kline, 2016). Root Mean Square Error of Approximation (RMSEA) was 0.024, which indicated excellent fit to the data (Hair, et al, 2010). Other indicators also proved the model fit, with CFI = 0.987, GFI = 0.938 and TLI = 0.986, all exceeded the instruction of 0.90 (Byrne, 2016).

**Table 9.** Structural Model Fit Indices.

| Fit Index | Value |
| --- | --- |
| CMIN/df | 1.235 |
| RMSEA | 0.024 |
| CFI | 0.987 |
| GFI | 0.938 |
| TLI | 0.986 |

#### b. Hypothesis Testing

The SEM findings indicated that all the hypothesized paths were statistically significant at the 0.001 level. Environmental Awareness (EA) and AI Engagement (AEEL) also had influential impacts on AT. Attitude, in return, Subjective Norms (SN) and Perceived Behavioral Control (PBC) had significate effects on Green Action Intention (GAI). Additionally, AEEL has positive significant influence on SN and PBC.

The direct determinant of GAI was PBC with the greatest standardized effect (β = 0.312), Attitude and SN were at second (β = 0.274) and third-ranked (β = 0.267) confirming their influence in determining the behavioral intention. Findings support the extended TPB model in the AI-powered environmental education setting.

**Table 10.** Regression Weights – Structural Paths.

| Hypothesized Path | Estimate ($\beta$) | S.E. | C.R. | p-value | Result |
| --- | --- | --- | --- | --- | --- |
| EA → AT | 0.236 | 0.052 | 4.538 | < .001 | Supported |
| AEEL → AT | 0.245 | 0.055 | 4.455 | < .001 | Supported |
| AEEL → SN | 0.281 | 0.058 | 4.847 | < .001 | Supported |
| AEEL → PBC | 0.252 | 0.057 | 4.421 | < .001 | Supported |
| AT → GAI | 0.274 | 0.059 | 4.644 | < .001 | Supported |
| SN → GAI | 0.267 | 0.063 | 4.238 | < .001 | Supported |
| PBC → GAI | 0.312 | 0.060 | 5.200 | < .001 | Supported |

All the structural model hypotheses were confirmed. The analysis supported the robustness of PBC, and then Attitude and SN, in predicting a green act intention among students. The model emphasizes the dual roles of psychological readiness and AI involvement in forming university students’ sustainable behaviors.

### 6. Mediation Analysis

Bootstrapped mediation with 5000 resamples in AMOS(NHTSA) To test the mediating role in our extended TPB model, we conducted the bootstrapping procedure with 5000 resampling the sample size. The mediation analysis was conducted to analyse whether AT, SN and PBC are the mediators of the relationships between the independent variables (EA, AEEL) and the dependent variable (GAI).

The indirect effects were found to be significant using the bootstrapping analysis. In particular, Attitude clearly mediated the association between EA and AEEL with GAI. Similarly, SN and PBC were the intermediates between AEEL and GAI. All indirect effects were statically significant (p < 0.05), with none of the confidence intervals containing zero, indicating the presence of substantive mediation.

In the majority of the instances, the indirect pathways were also accompanied by the presence of a significant direct path, indicating partial mediation. This shows that the effect of environmental awareness and AI engagement on green action intention has a direct role, as well as independent cognitive mediating roles.

**Table 11.** Bootstrap Results for Indirect Effects.

| Mediation Path | Indirect Effect | Boot SE | 95% CI | p-value | Mediation Type |
| --- | --- | --- | --- | --- | --- |
| EA → AT → GAI | 0.064 | 0.021 | [0.029, 0.108] | 0.002 | Partial |
| AEEL → AT → GAI | 0.067 | 0.023 | [0.030, 0.116] | 0.001 | Partial |
| AEEL → SN → GAI | 0.075 | 0.024 | [0.036, 0.127] | 0.001 | Partial |
| AEEL → PBC → GAI | 0.079 | 0.026 | [0.038, 0.134] | 0.001 | Partial |

Further, the mediation analysis supported that Attitude, Subjective Norms and Perceived Behavioral Control are key cognitive components through which AI engagement and environmental concern foster students’ intention for sustainable behavior. The observation of substantial partial mediation reveals the complex nature of behavioral shaping in the digital– environmental interplay.

### 7. Group Differences Analysis: T-Test and ANOVA

#### a. Gender Differences – Independent Samples T-Test

The Independent Samples T-Test was used for comparison of the males and females. Levene’s Test for Equality of Variances was performed before hypothesis testing. For all the variables the test showed that there was no significant violation of homogeneity (p > 0.05), enabling interpretation of equal variances assumed.

The findings indicate female students score higher than male students in terms of Environmental Awareness (EA), AI Engagement (AEEL) and Green Action Intention (GAI), and their p values are all below 0.05. This implies that female students are more interactively use environmental technologies and are more likely to participate in green behaviors, which is consistent with the trend in sustainability behavioral literature (Field, 2013).

**Table 12.** Independent Samples T-Test Results by Gender.

| Construct | Male Mean | Female Mean | t-value | p-value | Conclusion |
| --- | --- | --- | --- | --- | --- |
| Environmental Awareness (EA) | 3.78 | 4.12 | 2.867 | 0.004 | Significant |
| AI Engagement (AEEL) | 3.84 | 4.20 | 2.741 | 0.006 | Significant |
| Attitude (AT) | 4.15 | 4.22 | 0.985 | 0.326 | Not Significant |
| Subjective Norms (SN) | 3.95 | 4.03 | 1.102 | 0.272 | Not Significant |
| Perceived Behavioral Control (PBC) | 4.08 | 4.12 | 0.709 | 0.479 | Not Significant |
| Green Action Intention (GAI) | 3.76 | 4.05 | 2.631 | 0.009 | Significant |

#### b. Academic Year Differences – One-way ANOVA

Testing for school year-level differences One-Way ANOVA (analysis of variance) was employed to determine if the respondents’ school year-level (first year, second year, third year) significantly differs in their mean scores on the constructs of the model. Levene’s test was used to verify the expression of homogeneous variances (homogeneity of variances; p > 0.05). The ANOVA results found significant difference in AI Engagement, Subjective Norms and Green Action Intention (P< 0.05).

Tukey HSD post-hoc analysis showed that senior students reported significantly higher AI Engagement and Green Action Intention than freshmen. This trend may be a result of rising environmental content and use of digital tools over the years.

**Table 13.** One-Way ANOVA Results by Academic Year.

| Construct | F-value | p-value | Significant Difference Found |
| --- | --- | --- | --- |
| Environmental Awareness (EA) | 1.682 | 0.189 | No |
| AI Engagement (AEEL) | 4.205 | 0.016 | Yes (Final year > Year 1) |
| Attitude (AT) | 1.197 | 0.304 | No |
| Subjective Norms (SN) | 3.426 | 0.034 | Yes (Final year > Year 1) |
| Perceived Behavioral Control (PBC) | 1.803 | 0.169 | No |
| Green Action Intention (GAI) | 5.216 | 0.006 | Yes (Final year > Year 1) |

The findings both from T-Test and ANOVA show that demographic variables, such as gender and academic year, might have influences on some psychological and behavioral constructs in extended TPB model. There were a higher proportion of female and final year students who are environmentally literate, technologically active, and committed to environmental sustainability. The results of this study emphasize the need to design digital environmental education programs for diverse students to maximize the effect on their behavior.

## V. Implications

The research results have many theoretical, practical and policy applications, especially in the changing context of digital education and sustainable environment in Vietnamese higher education.

### 1. Theoretical implications

Developing TPB through adding factors as exogenous constructs to attitude, subjective norm and perceived control factors as they are backbones of the original TPB model can enable us not only to blossom the theory, but also to increase its application in different areas. With the confirmation of statistical significance that EA and AEEL have indirect effects on GAI through their mediators AT, SN, and PBC, the study supports the greater explanatory power of TPB in the digital and sustainability contexts.

In addition, the solid model fit indices of the structural model and the high reliabilities of construct indicate the robustness of this extended model and its legitimacy to apply to paralleled educational and cultural contexts. The substantial mediating effects of attitude and perceived control are consistent with current research in behavioral science, further supporting the relevance of the TPB to new domains (environmental informatics and eco-friendly technology in education).

### 2. Practical Implications for Educational Institutions

For universities (especially in climate-vulnerable areas like the Mekong Delta where this study was conducted), the study offers strong evidence that AI-driven digital platforms can be used to catalyze environmental awareness in students. Institutions such as Can Tho Private University that lead the use of smart learning technologies can make use of how engaged students are with AI in order to develop interactive, personalized and adaptive learning environments to infuse environmental content into their everyday teaching.

Additionally, due to the differences by gender and academic year on students’ responses, institutions should consider differentiated strategies. For example, fourth-year students demonstrated significantly more AI use and intention to behave sustainably that may relate to their learned knowledge and concerns about job security. This indicates that first year students need to be addressed with programs from the start which aim to provide a grounding in environmental literacy and digital skills. Likewise, gender-sensitive pedagogical interventions may increase participation and involvement for boys, who in this study showed lower green intentions and consciousness.

### 3. Policy and Curriculum Development

At a policy level, one implication is that building digital competencies and sustainability literacy into national education frameworks is an urgent requirement, not an optional choice. Ministries of Education and Training are encouraged to develop curricula that connect AI, environmental science, and behavioural education in interdisciplinary ways in order to prepare students to be critical, responsible, and competent digital citizens to tackle daunting, environmental grand challenges.

Furthermore, the findings call for greater cooperation among university authorities, local government, and nature protection societies to carry out university-wide green movements and computer-based instructional modules. These can be inscribed in international agendas like the UN Sustainable Development Goals (SDG 4, SDG 13, SDG 17).

### 4. Implications for Future Digital Sustainability Pedagogies

This work emphasizes the importance of reinterpreting sustainability education as not only a content-based, but also a behaviorally focused, digitally mediated learning process. The beneficial effect of AI exposure on the psychological antecedents of sustainability behaviour implies that carefully designed digital interventions (e.g., gamified eco-challenges, AI-based virtual labs or chatbot-supported environmental education) could possibly pave the way for a radical change in students’ habits and attitudes.

## VI. Conclusions

In the context of the rapidly increasing consequences of climate change occurring in the Mekong Delta, as well as the increasing reliance of artificial intelligence in re-moulding higher education, the current study explores the psychological and behavioral processes through which AI engagement and environmental awareness predict sustainable behavioral intention among university students.

The findings supported that both AI Engagement (AEEL), environmental awareness (EA) are direct and constructively related to Attitude (AT), Subjective Norms (SN), and Perceived Behavioral Control (PBC), as the significant dimensions in the TPB. In turn, these mediators had a significant impact on Green Action Intention (GAI) confirming the extended TPB model. Partial mediation was observed in the mediation analysis, which demonstrates that AEEL and EA have effects on behavioral intention not only directly, but also indirectly through cognitive-affective processes. The fit of the structural model was good, and all measurement constructs had high reliability and convergent validity.

The examination also found significant variance among demographic groups. Female students and students in their final semester obtained higher scores on environmental awareness, AI engagement, and intention to act in a sustainable manner. These variations imply that interventions in education should take students’ level of advancement into account in order to maximize behaviour outcomes.

Last, what this research finds as they come together is that digital transformation and environmental education can have a mutual synergy in influencing pro-environmental behavior. AI is not just a technology enabler, is a behavior amplifier and it is able to influence how students think, they feel and act towards environmental challenges. In the midst of Vietnam’s stride in national digitalization and green growth, universities should be the big brothers showing their reflections for a digital-citizenry upbringing and sustainable environment treatment.

In terms of the academic and practical implications, this study constitutes a significant perturbation from conventional theories and as such serves as a basis for rethinking how long-term behavior change with technology and sustainability could be combined in higher education.

## Notes

### Competing Interest Statement

The authors have declared no competing interest.

